# Contingency in the convergent evolution of a regulatory network: Dosage compensation in Drosophila

**DOI:** 10.1101/488569

**Authors:** Chris Ellison, Doris Bachtrog

## Abstract

The repeatability or predictability of evolution is a central question in evolutionary biology, and most often addressed in experimental evolution studies. Here, we infer how genetically heterogeneous natural systems acquire the same molecular changes, to address how genomic background affects adaptation in natural populations. In particular, we take advantage of independently formed neo-sex chromosomes in Drosophila species that have evolved dosage compensation by co-opting the dosage compensation (MSL) complex, to study the mutational paths that have led to the acquisition of 100s of novel binding sites for the MSL complex in different species. This complex recognizes a conserved 21-bp GA-rich sequence motif that is enriched on the X chromosome, and newly formed X chromosomes recruit the MSL complex by *de novo* acquisition of this binding motif. We identify recently formed sex chromosomes in the *Drosophila repleta* and *robusta* species groups by genome sequencing, and generate genomic occupancy maps of the MSL complex to infer the location of novel binding sites. We find that diverse mutational paths were utilized in each species to evolve 100s of *de novo* binding motifs along the neo-X, including expansions of microsatellites and transposable element insertions. However, the propensity to utilize a particular mutational path differs between independently formed X chromosomes, and appears to be contingent on genomic properties of that species, such as simple repeat or transposable element density. This establishes the “genomic environment” as an important determinant in predicting the outcome of evolutionary adaptations.

## Introduction

What would happen if we “replay the tape of life” (Gould 1989)? The question of whether adaptation follows a deterministic route largely prescribed by the environment or whether evolution is fundamentally unpredictable and can proceed along a large number of alternative trajectories has until recently been a fascinating problem that could not be addressed directly.

In the past decade, however, advances in DNA sequencing technology have allowed researchers to tackle this question using two complementary approaches. Experimental evolution of viruses, bacteria and yeast, in combination with genome sequencing, has allowed direct identification of adaptive mutations in order to address the relative contributions of determinism and stochasticity in the evolutionary process. Genomic analysis of populations experimentally evolved under controlled laboratory conditions has consistently revealed parallelism in which mutations in certain genes are repeatedly selected (Tenaillon *et al.* 2016; Venkataram *et al.* 2016). These studies are typically limited to systems that can be rapidly propagated in the lab and many relevant evolutionary parameters (such as environment, population size, etc.) are controlled by the experiment, and their applicability to natural systems is sometimes unclear (Barrick and Lenski 2013).

Studies of parallel adaptations in the wild are a complementary approach to understanding the repeatability of evolution. Organisms evolving under similar ecological conditions often evolve similar traits, and striking examples of genetic convergence at the DNA level have been recently discovered. For example, plant-feeding insects have independently and repeatedly colonized many different plant taxa, and highly diverged insect orders have evolved cardenolide resistance through the exact same amino acid substitution (Dobler *et al.* 2012). Similarly, three distantly-related lineages of snakes have convergently evolved resistance to the tetrodotoxin found in their prey via the same amino acid mutation in a voltage-gated sodium channel (Feldman *et al.* 2012). Phenotypic convergence can also result form noncoding changes. Parallel evolution of trichome patterning in Drosophila (Frankel *et al.* 2012) or wing pattern mimicry in Heliconius butterflies (Reed *et al.* 2011) both involved regulatory mutations that altered the expression pattern of a transcription factor. The parameters of convergent evolution in protein coding genes are fairly well understood and often involve a small number of amino acid mutations of large effect size that are constrained to specific regions of the protein because of pleiotropy. By contrast, how changes in *cis*-regulatory regions contribute to convergence is less well understood, and hampered by our limited understanding of the global cis-regulatory structure of a phenotype. Convergent regulatory evolution involves a much larger set of mutational targets and mechanisms: A single regulatory mutation affecting a transcription factor could act in trans to change the expression pattern of a suite of target genes (as observed in Drosophila and Heliconius), or multiple independent cis-acting mutations could act in concert to produce the selected phenotype. Furthermore, these regulatory mutations can arise via a large variety of mechanisms, from transposable element insertions to microsatellite expansions, and identifying causative adaptive mutations in nature is challenging, especially at noncoding DNA. Moreover, adaptation to novel habitats often involves multiple selective agents whose relative importance is often unclear, making interpretations of convergent evolution (or a lack thereof) challenging.

Here, we address how predictable evolution is at the DNA sequence level in nature, by studying the parallel evolution of a phenotype that is well understood at the molecular level: the acquisition of dosage compensation in fruit flies. Many species with heteromorphic sex chromosomes have evolved mechanisms to equalize the amount of gene product from the X chromosome in males and females (Vicoso and Bachtrog 2009). In Drosophila, males compensate for reduced dosage of X-linked genes by hyper-transcribing their hemizygous X chromosome through epigenetic modifications (Lucchesi 1978). At the molecular level, this is achieved by recruiting the male-specific lethal (MSL) complex to numerous chromatin entry sites (CES) on the X in a sequence-specific manner (Alekseyenko *et al.* 2008) (**Figure 1A**). The MSL complex targets a 21-bp long, GA-rich sequence motif that is enriched on the X chromosome (Alekseyenko *et al.* 2008). The complex then spreads from CES to the rest of the X, leading to chromosome-wide hyperactylation of H4K16, which results in upregulated transcription on the X chromosome (Lucchesi and Kuroda 2015).

**Figure 1.**
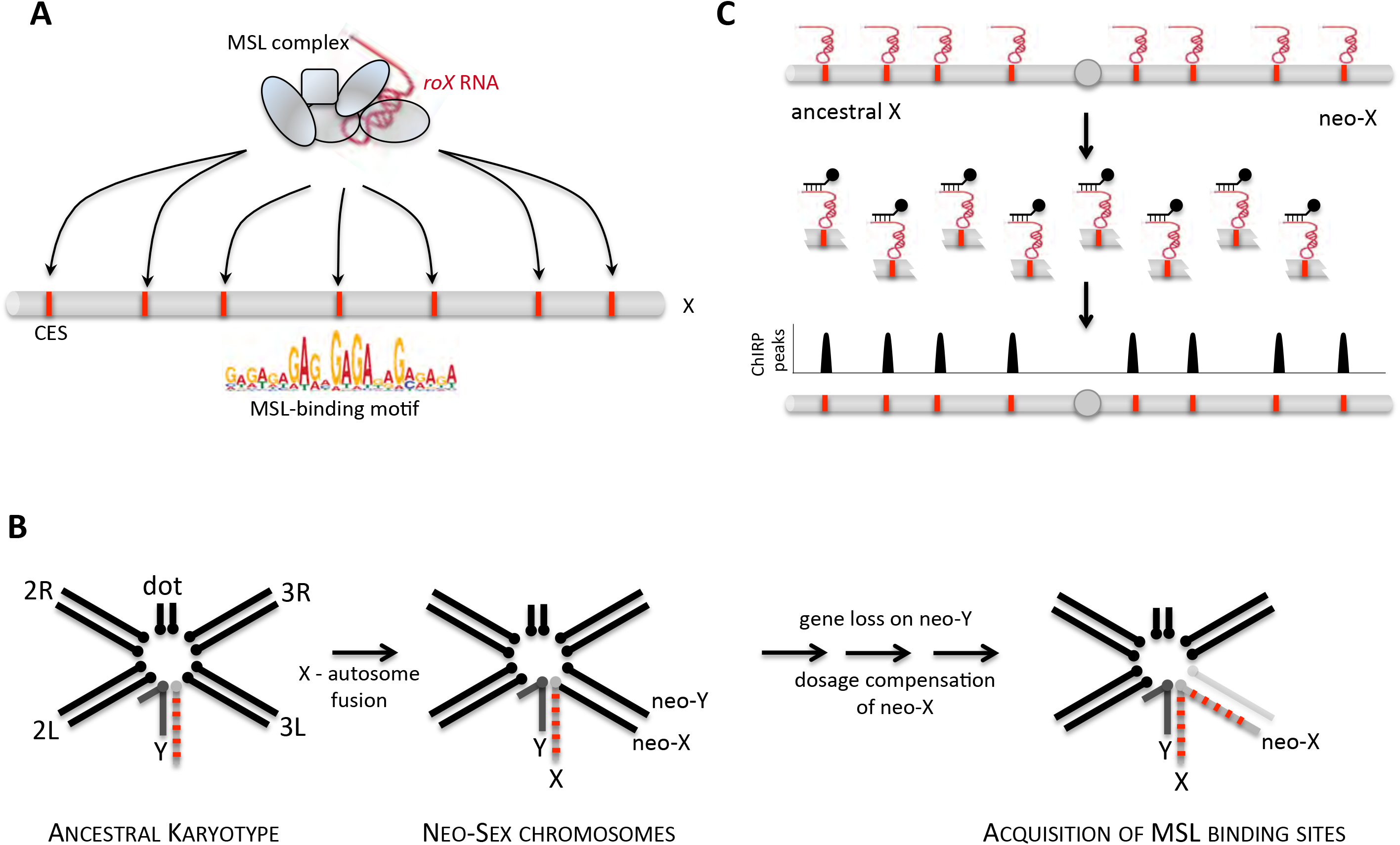
Dosage compensation and neo-sex chromosomes in Drosophila. **A.** MSL-mediated dosage compensation in Drosophila. The MSL-complex consists of several proteins and noncoding RNAs (*roX* RNA), and targets the X chromosome at chromatin entry sites (CES) that contain the MSL-binding motif (as GA-rich sequence motif). **B.** Formation of neo-sex chromosomes in Drosophila. The ancestral karyotype of Drosophila consists of 5 large rods (the ancestral X, which is conserved across Drosophila, and the autosomal arms 2L and 2R, and 3L and 3R) and the small dot chromosome. Autosomes repeatedly fused to the sex chromosomes, forming neo-X and neo-Y chromosomes. Loss of genes on the neo-Y creates selective pressure to dosage compensate neo-X genes, and has repeatedly evolved in Drosophila by co-opting the MSL-complex through the acquisition of novel MSL binding sites. **C.** ChIRP can be used to identify MSL-binding sites on Drosophila sex chromosomes. The *roX* RNA is bound *in vivo* to CES; chromatin is cross-linked and fragmented, and roX2 is affinity purified and sequenced.

MSL-mediated dosage compensation evolved over 60MY ago and is conserved across Drosophila species (Marín *et al.* 1996; Bone and Kuroda 1996). However, different species in this genus co-opted the MSL machinery to evolve dosage compensation on newly evolved neo-sex chromosomes (Marín *et al.* 1996; Bone and Kuroda 1996). In particular, fusions between the ancestral sex chromosomes (that is, the original X and Y chromosome shared by all members of the genus Drosophila) and autosomes have repeatedly created so-called neo-sex chromosomes (Muller). Once fused, the neo-sex chromosomes follow a distinct evolutionary trajectory over tens of millions of years until they obtain the classical properties of ancestral sex chromosomes: The neo-Y chromosome degenerates as its protein-coding genes are inactivated, and the neo-X chromosome is upregulated to compensate for this gene dosage imbalance (Lucchesi 1978; Bachtrog *et al.* 2008). During this transition, the age of the neo-sex chromosomes broadly correlates with their level of differentiation.

Dosage compensation evolves on newly formed X chromosomes by co-opting the MSL complex, through the acquisition of new MSL binding sites (**Figure 1B**). We recently studied the evolution of MSL binding sites in *D. miranda*, a model species for sex chromosome evolution that possesses two neo-X chromosomes that were formed about 13-15MY and 1.5MY ago, respectively (Bachtrog and Charlesworth 2002; Gao *et al.* 2007). We found that diverse mutational paths contributed to MSL binding site evolution (Zhou *et al.* 2013), but the majority of novel MSL sites on the young neo-X were created by insertions of a domesticated helitron transposable element (TE) containing the GA-rich sequence motif recognized by the MSL complex (the MSL recognition element, or MRE) (Ellison and Bachtrog 2013; 2015). We also detected highly eroded remnants of a related TE at the much older neo-X chromosome of this species, where dosage compensation evolved around 13-15 MY ago (Ellison and Bachtrog 2013).

Independently formed neo-X chromosomes are faced with the same evolutionary challenge: To co-opt the existing MSL machinery and up-regulate hundreds of genes simultaneously, in response to neo-Y degeneration. This creates a set of fascinating questions: How are new binding sites acquired on different positions along the neo-X chromosome of a lineage, or on independently evolved neo-X chromosomes between species? Does evolution predominantly follow the same molecular path to evolve new MSL binding sites, do species-specific solutions evolve to the same problem, or do independent binding sites evolve by diverse molecular mechanisms even within a lineage?

To address these questions, we use comparative genomic and functional analysis to infer the mutational path evolution has taken to acquire novel MSL binding sites. We focus on Drosophila species in the *melanica* and *robusta* groups, a promising system to study the independent rewiring of the MSL complex. Cytogenetic studies have shown that species in this clade have independently evolved neo-sex chromosomes, thus ensuring that dosage compensation evolved in parallel for these species. Phylogenetic dating suggests that neo-sex chromosomes in this group are young (Flores *et al.* 2008), which should allow us to identify the causative mutations that created novel MSL binding sites on neo-X chromosomes.

In this study, we generate genomic data for five species from the *melanica*/*robusta* groups, to identify the specific X-autosome fusions in these species and date their formation. We create maps of MSL occupancy for three species where dosage compensation on the neo-X has evolved independently and recently. Comparative analysis allows us to reconstruct the mutations generating novel MSL binding sites, and we infer both heterogeneity and convergence of binding site evolution within and between species. Our results demonstrate that evolution is highly opportunistic, yet contingent on the genomic background. We show that species use a diverse spectrum of mutational events to generate novel MSL binding sites, but the propensity for different types depends on genomic contingencies of a species.

## Results

### Identification of independently formed neo-sex chromosomes in Drosophila

Young neo-sex chromosomes of Drosophila have formed independently by fusions between autosomes and the ancestral sex chromosomes (Lucchesi 1978). Over time, they acquire the stereotypical properties of the ancestral X and Y, and their repeated formation in different lineages at different time points allows us to contrast neo-sex chromosomes at various stages of differentiation (Zhou and Bachtrog 2015; Mahajan *et al.* 2018). Cytogenetic comparisons suggest that neo-sex chromosomes have evolved independently multiple times in species from the *virilis-repleta* radiation (Flores *et al.* 2008), but their neo-sex chromosomes were not examined at the genomic level. To identify neo-sex chromosomes and infer their evolutionary history, we performed Illumina whole-genome sequencing of males and females from five species in the *robusta/melanica* sister groups (**Figure 2; Table S1**). We generated *de novo* assemblies from the female sequencing data and created a whole-genome alignment to identify orthologous regions between all five species, which we used to infer their phylogeny. Our genome-wide phylogenetic analysis confirms previous inferred relationships among members of these groups based on a handful of genes (Wang *et al.* 2006; Flores *et al.* 2008), with *D. melanica* and *D. nigromelanica* being sister species, and *D. micromelanica* as their outgroup (forming the *melanica* group), whereas *D. robusta* and *D. lacertosa* are more distantly related (**Figure 2A**). We used sequence divergence between species pairs to roughly date their split times. Assuming a neutral mutation rate of 3.46 x 10^-9^ per year (Keightley *et al.* 2009), we estimate that species from the *melanica* subgroup split very recently, *D. melanica* and *D. nigromelanica* diverged roughly 4.3 MY ago, *D. robusta* split from the *melanica* species about 9.4 MYA, and *D. lacertosa* diverged 16 MY ago (**Figure 2B**).

**Figure 2.**
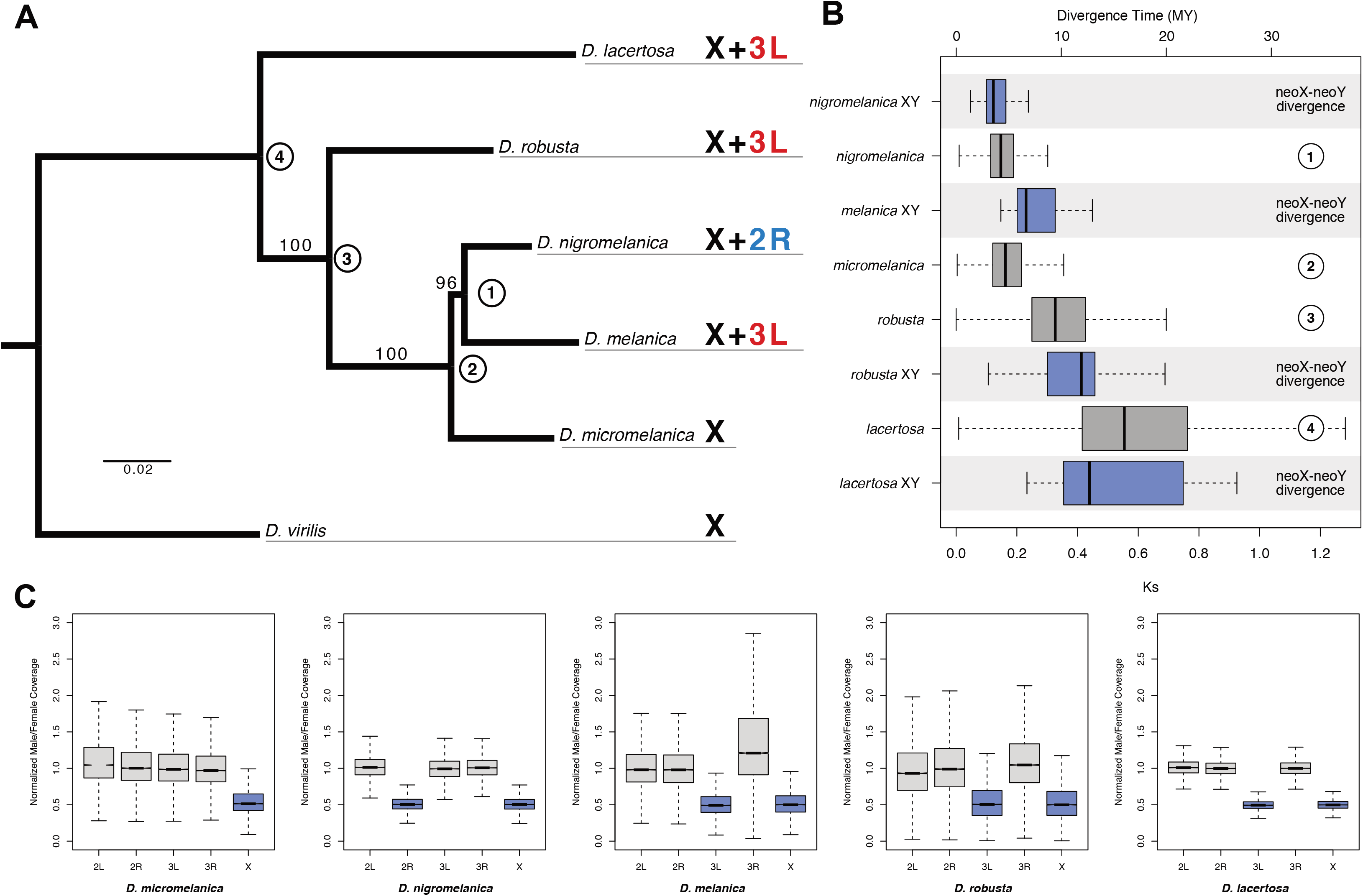
Phylogenetic relationships and karyotype evolution in the *melanica* and robusta species groups. A. Whole-genome maximum likelihood phylogeny inferred using the RAxML rapid bootstrapping algorithm. Bootstrap support values are shown on branches and nodes are numbered for reference to Panel C. Each species is labeled according to which chromosomes are X-linked. X refers to the ancestral X chromosome, while 3L and 2R refer to the *D. melanogaster* autosomes Muller D and Muller C. B. Divergence (calculated as Ks and converted to MYa) between neo-X/Y chromosomes as well as the nodes labeled in Panel A. Boxes reflect the distribution of Ks values for all neo-X/Y gene pairs or between-species orthologs. C. X-linkage was determined based on the ratio of Male/Female Illumina sequencing coverage for genomic libraries aligned to the female assemblies (autosomes=grey, X chromosomes=blue).

Neo-sex chromosomes were previously reported for four of the five species investigated here, with *D. micromelanica* lacking an X-autosome fusion. The neo-sex chromosomes of *D. nigromelanica* and *D. melanica* were thought to be homologous (Flores *et al.* 2008), while X-autosome fusions occurred independently in *D. robusta* and *D. lacertosa* (Flores *et al.* 2008). We used male and female genomic coverage data to infer which autosomal chromosome arm became the neo-sex chromosome in each species, and to estimate the date at which the fusion occurred (see Methods). As expected, we identified the ancestral X chromosome (which is shared across Drosophila) by reduced male coverage in each species, and we confirmed the presence of a neo-X chromosome in *D. nigromelanica, D. melanica, D. robusta and D. lacertosa*, as well as the absence of a neo-X in *D. micromelanica* (**Figure 2C**). Intriguingly, however, we find that a different chromosome arm formed the neo-sex chromosome in the *melanica* group: chromosome 3L became the neo-sex chromosome of *D. melanica*, while chromosome 2R formed the neo-sex chromosome of *D. nigromelanica*. Thus, contrary to parsimonious interpretations based on cytological data, our genomic comparison shows that neo-sex chromosomes formed independently at least two times in the *melanica* group and imply that they are younger than previously assumed. We also confirmed that chromosome 3L became the neo-sex chromosome of both *D. lacertosa* and *D. robusta* (**Figure 2C**), but the lack of X-autosome fusions in multiple species of the *lacertosa* and *robusta* subgroups indicates that these fusions originated independently (Flores *et al.* 2008).

The age of each species group with unique or shared neo-sex chromosomes sets a limit to the age of their chromosomal fusions (**Figure 2B**). Phylogenetic analysis suggests that *D. melanica* and *D. nigromelanica* diverged about 4.3 MY ago, implying that both species groups’ neo-sex chromosomes are younger than that age. Sequence divergence of homologous neo-sex linked genes provides an independent estimate of their age: older neo-Y chromosomes harbor fewer genes (and fewer neo-X/neo-Y gene pairs), and levels of sequence divergence between orthologous gene pairs increases with the age of the neo-sex chromosome. We identified 118 pairs of homologous neo-sex linked genes in *D. nigromelanica*, 20 in *D. melanica*, 16 in *D. robusta* and 11 in *D. lacertosa*, and sequence divergence between homologous gene pairs increases with decreasing gene number (**Table S2, Figure 2B**). This analysis suggests that *D. nigromelanica* has a relatively young neo-sex chromosome (mean dS=0.16, 4.6 MY), followed by *D. melanica* (mean dS=0.26, 7.5 MY), while the sex chromosomes of *D. robusta* and *D. lacertosa* formed about 11-15 MY ago mean dS=0.39, 0.52; 11.3, 15.0 MY). The inferred ages of neo-sex chromosomes are in between those for the neo-sex chromosomes of *D. miranda*, whose older neo-sex chromosome (chromosome XR) fused to the ancestral X about 13-15MY ago, and its neo-X/neo-Y formed about 1.5MY ago (Bachtrog and Charlesworth 2002). Note that the estimated age of neo-sex chromosomes may exceed the inferred speciation time even if they formed after speciation due to faster sequence evolution on the neo-Y chromosome. Selection is less efficient on the non-recombining neo-Y chromosome (Bachtrog and Charlesworth 2002) and slightly deleterious synonymous mutations may thus accumulate faster and a higher mutation rate in males relative to females (male-driven evolution) could further increase the rate of neutral substitutions along the neo-Y branch (Bachtrog 2008).

### Neo-X chromosomes have acquired dosage compensation

Degeneration of the Y chromosome creates selective pressure to dosage compensate the X chromosome (Bachtrog 2013). Our genomic analysis demonstrates that the neo-Y chromosomes of members of the *robusta* and *melanica* species group have very few genes left, and their neo-X chromosomes may thus have already acquired full dosage compensation. We gathered male and female RNA-seq data from *D. melanica* and *D. robusta* heads, to test if expression levels of the newly formed neo-X chromosomes are similar between males and females (i.e. whether they have evolved dosage compensation). We assigned *melanica* and *robusta* genomic scaffolds to chromosomes based on homology to *D. virilis* and identified genes directly from RNA-seq alignments to those same genomic scaffolds. **Figure 3A** shows male/female expression ratios for genes on the different chromosomes. Genes located on the neo-X in both species show very similar male/female expression ratios to genes on the ancestral X, suggesting that they have evolved full dosage compensation. Interestingly, however, genes located on the ancestral X as well as the neo-X in both species show slightly higher expression in females than males, compared to autosomes (**Figure 3A**). This is consistent with the female-biased expression pattern of X chromosomes previously observed in Drosophila (Sturgill *et al.* 2007; Assis *et al.* 2012).

**Figure 3.**
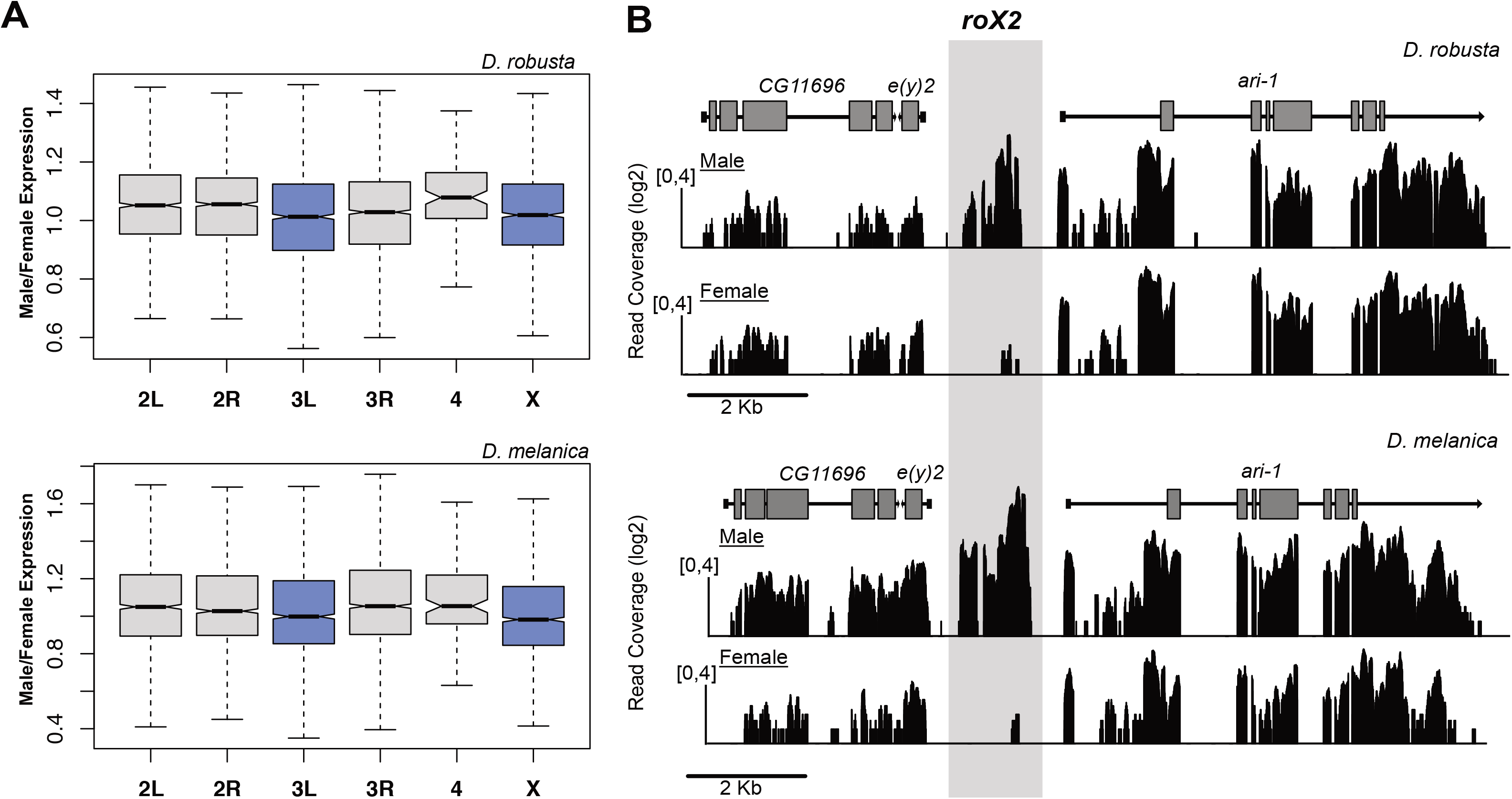
Dosage compensation of neo-X chromosomes. A. We profiled gene expression in Drosophila heads using RNA-seq and Male/Female gene expression ratios are shown for each chromosome for *D. robusta* (top) and *D. melanica* (bottom). X-linked genes (blue boxes) are expressed at roughly equal levels in both Males and Females, suggesting that both the ancestral and neo-X chromosomes are fully dosage compensated. **B.** Identification of roX2. In other Drosophila species, the male-specific non-coding roX2 transcript is located between the protein-coding genes *e(y)2* and *ari-1*. We identified the roX2 transcript in *D. melanica* and *D. robusta* by visualizing RNA-seq data from heads across the *e(y)2* – *ari-1* genomic region. The grey box indicates the location of roX2, which shows abundant expression only in males.

### Annotation of *roX* sequences and identification of MSL-binding sites

Comparative studies in Drosophila have shown that dosage compensation by the MSL complex is conserved across species (Marín *et al.* 1996; Bone and Kuroda 1996; Alekseyenko *et al.* 2013; Quinn *et al.* 2016). Moreover, newly formed neo-X chromosomes evolve dosage compensation by acquiring novel MSL-binding sites which are able to recruit the MSL complex (Alekseyenko *et al.* 2013; Quinn *et al.* 2016). In *D. melanogaster*, two non-coding RNAs are part of the MSL-complex, and a recently developed technique known as Chromatin Isolation by RNA Purification, or ChIRP, has been successfully used to map MSL binding in several Drosophila species, by isolating and sequencing DNA bound by the *roX* non-coding RNAs (Quinn *et al.* 2016) (see **Figure 1C**).

Non-coding RNAs evolve quickly at the DNA sequence level, but can be identified based on microsynteny, and their male-specific expression (Quinn *et al.* 2016). Previous work has shown that while *roX1* strongly localizes to the X chromosome in *D. melanogaster*, it shows much weaker X-localization in other species of Drosophila (including *D. virilis*) (Quinn *et al.* 2016). *RoX2*, on the other hand, shows strong localization to the X chromosome in all Drosophila species studied so far (Quinn *et al.* 2016), and has male-specific expression in dozens of species across the Drosophila phylogeny (Quinn *et al.* 2016); we thus focused on identifying *roX2* in our target species. In *D. virilis*, *roX2* is located between the protein-coding genes *ari-1* and *e(y)2*. To identify the *roX2* locus, we first searched each genome for synteny blocks likely containing *roX2* based on the conserved location of *ari-1* and *e(y)2* homologs, and then mapped RNA-seq data from *D. melanica* and *D. robusta* males and females, in order to identify genomic regions showing male-specific expression (**Figure 3B**). Indeed, we found male-specific RNA-seq reads from our candidate region in both species, and we assembled the RNA-seq reads mapping to this location to generate the full-length *roX2* transcripts from each species. We identified *roX2* in *D. nigromelanica* based on homology to the *D. melanica* and *D. robusta* transcripts (**Figure S1**).

To map MSL-binding sites, we designed non-overlapping oligos against *roX2* in *D. nigromelanica*, *D. melanica*, and *D. robusta*, using a split oligo design (Quinn *et al.* 2016). We performed two independent ChIRP experiments with different non-overlapping oligo sets (**Table S3**), and generated 100-bp paired-end sequencing libraries for each oligo set as well as an input control. We aligned the sequencing reads to their respective genomes, and identified genomic regions showing *roX2* enrichment peaks. We found that *roX2* binding is highly correlated between independent probe set for each of the species **(Figure S2**), and we identified MSL-binding sites as overlapping peaks between the two independent ChIRP experiments. We identified between 980 and 1,570 peaks in each species, with the majority located on either the X or neo-X chromosome (~75-95%, **Figure 4A**). As done previously (Alekseyenko *et al.* 2013; Quinn *et al.* 2016), we set an enrichment threshold (see Methods) to identify the subset of the most strongly bound peaks as chromatin entry sites (CES). We found similar numbers of CES on the ancestral X (212-295) and the neo-X (258-290), suggesting that the neo-X chromosomes in each species have evolved full dosage compensation. This is consistent with gene expression patterns, and the number of CES per chromosome are similar to what has been found in other species with fully dosage compensated X chromosomes (Alekseyenko *et al.* 2008).

**Figure 4.**
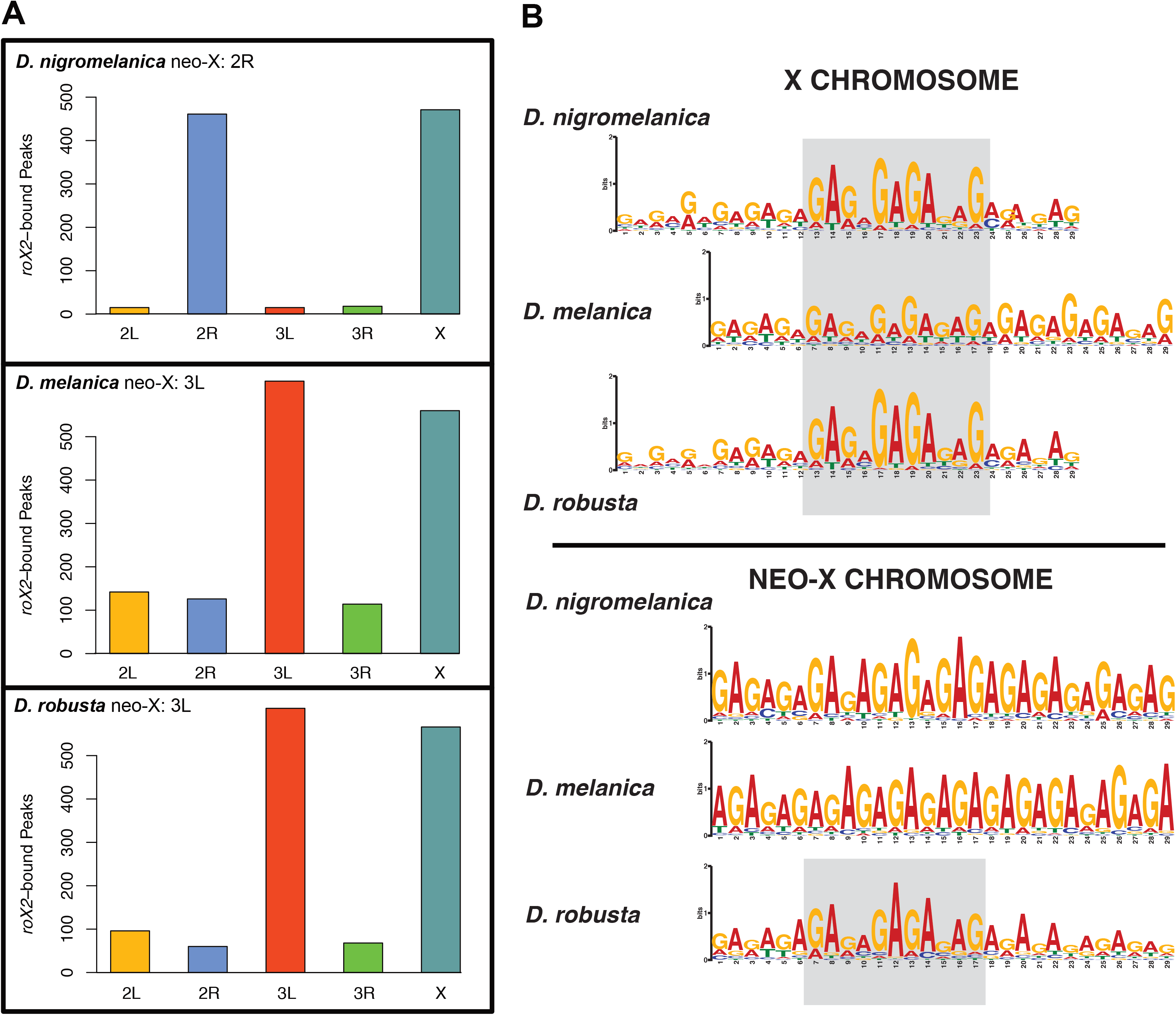
*roX2* binding locations and sequence motifs. A. *roX2* bound regions are highly enriched on both the ancestral X chromosome and the neo-X chromosome in all three species. B. Identification of MSL binding motif. From the strongly bound peaks corresponding to chromatin entry sites (CES), we identified a GA-rich sequence motif on both the ancestral and neo-X chromosomes in all three species. The grey boxes highlight the conserved core motif, which is very similar to that found in *D. melanogaster*.

We used MEME to identify motifs enriched within *roX2*-bound CES regions. Careful molecular studies in *D. melanogaster* identified a GA-rich sequence motif that is targeted by the MSL-complex (Alekseyenko *et al.* 2013). Consistent with previous studies, we find a GA-rich sequence motif to be highly enriched on the ancestral X in every species studied (**Figure 4B**), and the same GA-rich sequence motif is found on the newly evolved X chromosomes in species of the *robusta* and *melanica* species group. A subset of CES, called pioneering sites on the X (pionX sites) are thought to be responsible for the initial recruitment of the MSL complex to the X chromosome (Villa *et al.* 2016). PionX sites share the low-complexity GA-rich motif with canonical CES, but contain a more complex CAC 5’ extension (Villa *et al.* 2016). We searched each CES to identify matches to both the canonical MRE motif and the pion-X motif (**Figure S3**). Between 85-91% of the CES sites identified contain either a MRE or pion-X motif on both the ancestral X and on the neo-X (174/175/139 MRE and 84/65/57 pion-X on the ancestral X and 137/169/157 MRE and 73/82/86 pion-X on the neo-X in *D. nigromelanica* / *D. melanica* / *D. robusta*; **Table 1**). Thus, this suggests that the molecular machinery for dosage compensation is conserved in Drosophila, and that the MSL complex has been independently recruited to transcriptionally up-regulate the newly formed X chromosomes by acquisition of the MSL binding motif.

**Table 1.** CES identified in *D. nigromelanica*, *D. melanica* and *D. robusta*.

|  | X chromosome |  |  | Neo-X chromosome |  |  |
| --- | --- | --- | --- | --- | --- | --- |
|  | MRE | Pion-X | No motif | MRE | Pion-X | No Motif |
| *D. nigromelanica* | 174 (59%) | 84 (27%) | 37 (13%) | 137 (53%) | 73 (28%) | 48 (19%) |
| *D. melanica* | 175 (64%) | 65 (24%) | 35 (13%) | 169 (58%) | 82 (28%) | 39 (13%) |
| *D. robusta* | 139 (66%) | 57 (27%) | 16 (8%) | 157 (59%) | 86 (32%) | 25 (9%) |

### CES conservation and turnover on the ancestral X

Contrasting CES evolution on homologous chromosomes that either ancestrally or convergently evolved dosage compensation allows us to study evolutionary patterns and constraints of CES conservation, acquisition, and turnover (Alekseyenko *et al.* 2013; Zhou *et al.* 2013; Ellison and Bachtrog 2013; Quinn *et al.* 2016). In particular, the ancestral X was fully dosage compensated by the MSL complex in the last common ancestor of *D. melanica*, *D. nigromelanica* and *D. robusta*, and contrasting CES locations on the ancestral X chromosome can inform us on the evolutionary stability and turnover of shared CES. In contrast, chromosome 3L independently evolved dosage compensation in *D. melanica* and *D. robusta,* and convergent acquisition of CES on homologous positions may reflect either the presence of dosage-sensitive genes in a particular location and strong need to evolve CES, or the presence of pre-sites (i.e. nucleotide sequences that resemble the MRE motif) and thus an easy mutational path to acquire CES.

Overall, we find 72 CES (about 28%) and 46 motifs (20%) to be syntenic between all three species on the ancestral X chromosome, and 50% of CES sites (and 41% of motifs) are shared between the more closely related *D. melanica* and *D. nigromelanica* species pair (**Table S4**). Each species from the *melanica* subgroup shares about 43% of its CES sites with the more distantly related *D. robusta*. Inspection of species-specific CES reveals that the majority of orthologous regions in other species (77% on average) tend to be bound by roX2, but at too low a level to pass our genome-wide threshold for CES identification; the majority of orthlogous regions also contain the MRE/pionX motif (73% on average), suggesting that most CES on the ancestral X are conserved between species.

On the other hand, about 25% of CES evolved independently at syntenic positions on chromosome 3L in *D. melanica* and *D. robusta* (i.e. 54 out of 251/243), suggesting that they arose from a presite present in their common ancestor. Indeed, for 90% of these sites, the orthologous region in *D. nigromelanica*, where this chromosome is an autosome, shows homology to the MRE/pion-X motif, indicating that the MSL binding site evolved from a pre-existing pre-site. Five genomic regions evolved CES independently in *D. melanica* and *D. robusta*, either by GA-expansions, or insertions.

### Mutational paths to acquire MSL binding sites on the neo-X

How are novel CES acquired at the molecular level? Previous analysis of CES evolution in Drosophila revealed diverse mutational paths by which novel CES originated in different fly species (Zhou *et al.* 2013; Ellison and Bachtrog 2013; Quinn *et al.* 2016). They include the use of pre-binding sites (i.e. sequence motifs that resemble the MRE motif and predate CES formation), simple GA expansions (Kuzu *et al.* 2016), or the spreading of CES by TE mobilization (Ellison and Bachtrog 2013). Overall, we find that about 40-60% of CES on the newly formed neo-X chromosomes evolved from a pre-site. Thus, CES often evolve from sequences that ancestrally resemble the MRE or pion-X motif, and changes to either the flanking sequences of CES (Alekseyenko *et al.* 2012), the repeat composition of the X (Joshi and Meller 2017) or possibly changes to the 3D organization (Schauer *et al.* 2017) may allow these sequences not previously targeted by the MSL complex to function as CES. In *D. melanica* and *D. nigromelanica*, most remaining CES (~25 %) are created by simple GA expansions, whereas in *D. robusta*, 28% of novel CES were created by transposable element insertions (**Figure 5A, 5B**). Thus, all three species utilized diverse mutational paths to evolve 100s of novel CES on their independently formed neo-X chromosome.

**Figure 5.**
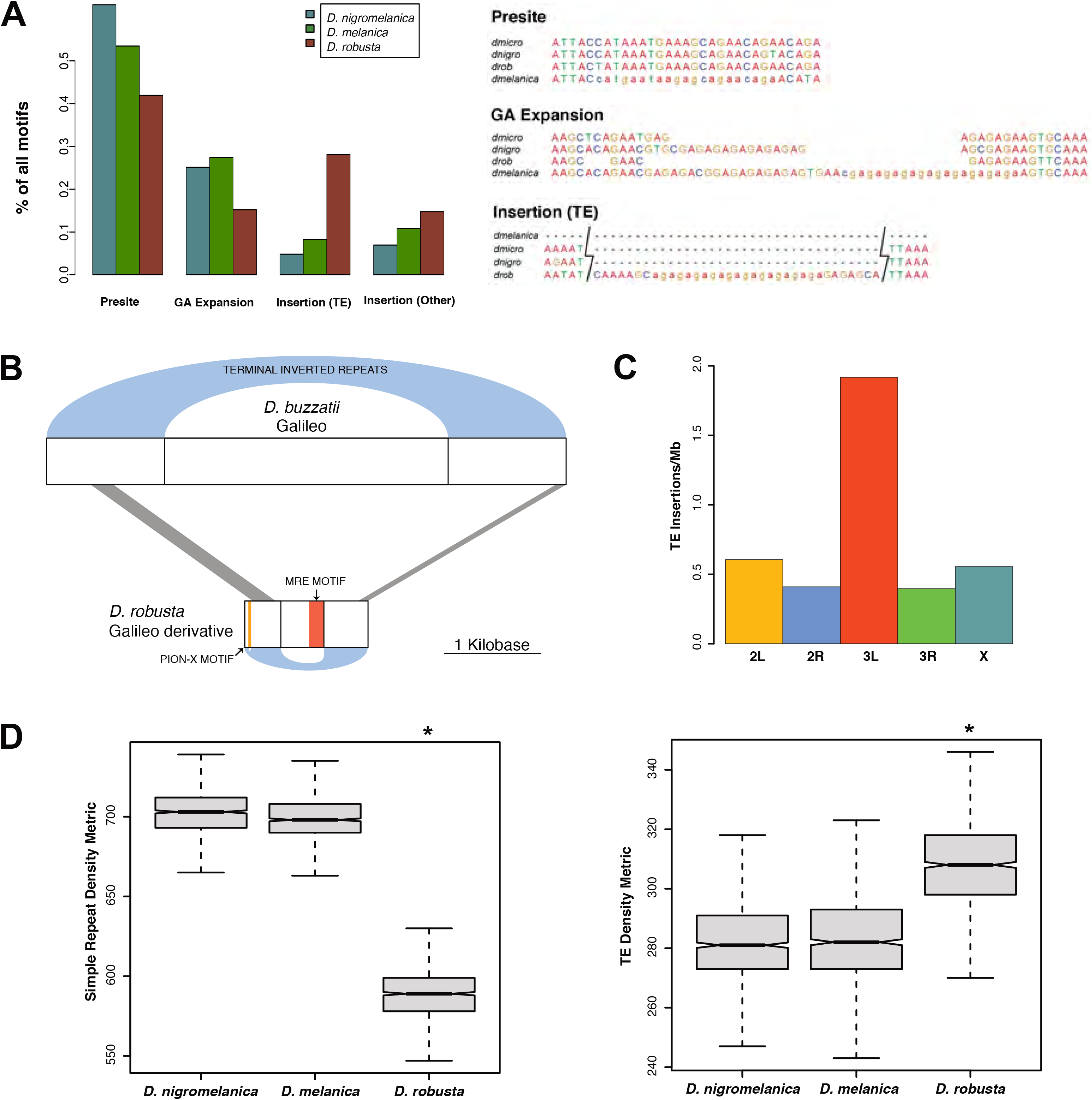
Evolution of chromatin entry site (CES) motifs. A. We used a comparative genomics approach to determine the evolutionary mechanism that gave rise to the MSL-binding motifs found within each CES on the neo-X chromosome of each species; shown is one example of each major type. *D. robusta* has a larger number of CES motifs derived from transposable element insertions, compared to the other two species, whereas *D. nigromelanica* and *D. melanica* have a larger number of motifs that arose from GA-expansion. B. We identified a Galileo-like TE that gave rise to at least 24 CES motifs on the *robusta* neo-X chromosome. C. Insertions of this TE are significantly enriched on the *robusta* neo-X, compared to both autosomes and the ancestral X chromosome (Binomial Test *P*=3.2e-14). D. Both *D. nigromelanica* and *D. melanica* genome assemblies have a higher density of simple repeats, whereas the *D. robusta* genome assembly has a higher density of TEs (Wilcoxon Test P<2.2e-16 for both comparisons). The repeat density metric refers to the number 1 kb windows that overlap a repeat in the genome assembly out of 1000 randomly placed windows.

We also compared the mechanisms that gave rise to MRE motifs versus those that produced pion-X motifs separately (**Figure S4**). The relative frequencies of each mutational mechanism were similar for both types of motifs, with the exception that MRE motifs were much more likely to arise from GA-expansions compared to pionX motifs (Fisher’s Exact Test *P*=4.9e-08), consistent with the more complex sequence content of the pion-X motif. A recent study suggested the CES might originate by the co-option of GA-rich polypyrimidine tracts that are located at the 3’ 100bp of introns and are used for splicing (Quinn *et al.* 2016)). We found that 7% - 12% of neo-X CES motifs are within the 3’ 100bp of introns, the expected location for motifs arising from a co-opted polypyrimidine tract. Both MRE and pionX motifs arose from these locations (55%-76% MRE, 24%-45% pion-X) and in roughly half of these cases, the polypyrimidine tract served as a ready-made motif (i.e. presite), whereas in the remaining half, there were additional GA-expansion mutations, suggesting that the co-option of these features can involve multiple mutational paths (**Figure S5**).

We investigated the *D. robusta* TE-derived motifs in more detail and found that both MRE and pionX motifs were derived from a total of 18 different families (**Table S5**). While most of these TE families were associated with only a single motif, we also identified a single family of elements that gave rise to 24 MSL binding motifs (17 MRE motifs and 7 pionX motifs). This TE family contains terminal inverted repeats with weak homology to those of two related *galileo* elements that have been identified in *D. buzzatii* and *D. melanogaster* (**Figure 5B**). The TE consensus sequence contains a match to the pionX motif near its 5’ end as well as a MRE motif near its 3’ end. We identified several hundred fragmented copies of this TE in our *D. robusta* genome assembly. These copies are highly enriched on the *D. robusta* neo-X chromosome (Binomial Test *P*=3.2e-14, **Figure 5C**) and overlap CES more often than expected by chance (Fisher’s Exact Test *P*<2.2e-16). Of the 66 copies that are located on scaffolds that were assigned to Muller elements, 32 are from the neo-X chromosome and 24 of these 32 copies lie within CES. The average pairwise genetic distance between these copies is 0.47, which suggests they were active around the time when the neo-sex chromosomes of *D. robusta* were formed (mean dS: 0.39). Thus, enrichment of this TE on the neo-X of *D. robusta*, its occurrence within CES and the presence of a binding motif for the MSL complex, and its inferred time of mobilization around the formation of neo-sex chromosomes strongly suggests that this TE was actively involved in dispersing MSL-binding sites along the neo-X.

### Genomic contingencies and heterogeneity in binding site evolution

What drives heterogeneity in CES acquisition across lineages? While *D. melanica* and *D. nigromelanica* evolved most novel CES by GA expansions, *D. robusta* utilized a TE for acquiring novel CES. To investigate if certain genomic factors prime a genome to preferentially evolve CES by a particular mutational path, we analyzed the content of repetitive DNA – both microsatellite and TE density, in the different lineages. Interestingly, we found that *D. melanica* and *D. nigromelanica* differ in their overall repeat composition from *D. robusta*: they both show a higher density of simple repeats but a lower density of TE sequences (Wilcoxon Test *P*<2.2e-16 for both comparisons)(**Figure 5D**). Higher TE density in *D. robusta* is consistent with its larger assembled genome size (185 Mb in *D. robusta* vs. 150 Mb in *D. melanica* and 164 Mb in *D. nigromelanica*). Thus, this observation is consistent with the notion that historical contingencies constrain evolutionary patterns of MSL binding sites evolution. TE’s may be more often utilized for rewiring regulatory networks in species with a higher number of TEs, but a larger TE burden may also contribute to increased genome sizes. On the other hand, a higher density of simple satellites in D. melanica and *D. nigromelanica* may have pre-adapted them to evolve novel MRE sites by GA-sequence expansion.

## Discussion

We took advantage of naturally occurring variation in sex chromosome karyotype in Drosophila species, to study independent replicates of solving the same evolutionary challenge: to dosage compensate newly formed neo-X chromosomes by acquiring hundreds of MSL-binding sites, in response to Y degeneration.

The independent acquisition of dosage compensation in Drosophila allows us to address several important questions in evolutionary biology and gene regulation: First, how repeatable is evolution? Evolutionary biologists have long debated the predictability of the evolutionary process. At one extreme, evolution could be highly idiosyncratic and unpredictable, since the survival of the fittest could occur along a great number of forking paths. Alternatively, constraints on evolution may force independent lineages to frequently converge on the same genetic solutions for the same evolutionary challenge. Second, how do regulatory networks evolve? And what is the contribution of TEs to regulatory evolution? Evolutionary innovations and adaptations often require rapid and concerted changes in regulation of gene expression at many loci (Carroll 2006). Transposable elements constitute the most dynamic part of eukaryotic genomes, and the dispersal of TEs that contain a regulatory element may allow for the same regulatory motif to be recruited at many genomic locations, thereby drawing multiple genes into the same regulatory network (McCLINTOCK 1951; Britten and Davidson 1969; Rebollo *et al.* 2012). Third, what makes a binding motif functional? The genomes of complex organisms encompass megabases of DNA, and regulatory molecules must distinguish specific targets within this vast landscape. Regulatory factors typically identify their targets through sequence-specific interactions with the underlying DNA, but they typically bind only a fraction of the candidate genomic regions containing their specific target sequence motif. An unresolved mystery in regulatory evolution is what drives the specificity of binding to a subset of genomic regions, that all appear to have a sequence that matches the consensus binding motif.

Several features make dosage compensation in Drosophila a promising system to tackle these questions. The genetic architecture for most adaptations, especially those involving regulatory changes, as well as the timing and exact selective forces driving them are generally little understood. In contrast, we have detailed knowledge of the molecular mechanism of dosage compensation in Drosophila. We know the cis- and trans-acting components of this regulatory network, and the regulatory motif for targeting the MSL complex to the X. We have clear expectations of which genomic regions should acquire dosage compensation, and about the timing and the evolutionary forces that drive wiring of hundreds of genes into the dosage compensation network on newly evolved X chromosomes. Specifically, Y degeneration is a general facet of sex chromosome evolution, creating selective pressures to up-regulate X-linked genes in males. Dosage compensation should thus only evolve on neo-X chromosomes whose neo-Y homologs have started to degenerate, and should evolve simultaneously or shortly after substantial gene loss has occurred on the neo-Y. Thus, our refined understanding of the how, when, why and where of dosage compensation in Drosophila makes this an ideal model system to study the repeatability of evolution, and the evolution of regulatory networks.

### How predictable is evolution?

Results from evolution experiments indicate that although evolution is not identical in replicate populations, there is an important degree of predictability (Lässig *et al.* 2017). Experimentally evolved populations under controlled, identical conditions consistently show parallelism in which mutations in certain genes are repeatedly selected (Tenaillon *et al.* 2016; Venkataram *et al.* 2016). However, organisms adapting to similar environments are not genetically identical, but their genome instead carries the legacy of their unique evolutionary trajectory, raising the question of how genomic differences affect genetic parallelism.

Sex chromosome - autosome fusions have independently created neo-sex chromosomes in different Drosophila lineages. This provides us with several independent replicates to study how, on the molecular level, evolution has solved the same adaptive challenge: acquiring hundreds of binding sites to recruit the MSL complex to newly formed X chromosomes. This allows us to quantify how much variation there is, both within and between species, in the underlying mutational paths to acquire hundreds of MSL-binding sites on neo-X chromosomes and identify genomic contingencies that will influence the repeatability of evolutionary trajectories. Importantly, neo-sex chromosomes of Drosophila are evolutionary young (between 0.1-15 MY old), which allows us, in many cases, to infer the causative mutations that have resulted in the gain of a regulatory element, and decipher the evolutionary processes at work to draw hundreds of genes into a new regulatory network.

Our results suggest that evolution is highly opportunistic, but contingent on genomic background. In particular, we find that each independently evolved neo-X chromosome uses a diverse set of mutational pathways to acquire MSL binding sites on a new neo-X chromosome, ranging from microsatellite expansions, the utilization of pre-sites, or transposable element insertions. However, different lineages differ with regards to the frequency of which mutational paths are most often followed to acquire novel binding sites, and this propensity may depend on the genomic background. In particular, we find that the two species with the higher density of simple repeats are more prone to utilize expansions in GA-microsatellites to gain a novel MSL binding sites. In contrast, *D. robusta* has an elevated TE density compared to its sibling species, and we find that the dispersal of a TE has played an important role in the acquisition of MSL binding sites on its neo-X chromosome. Thus, this suggest that the genomic background of a species predisposes it to evolve along a particular path, yet the evolutionary process is random and resourceful with regards to utilizing a variety of mutations to create novel MSL binding sites.

### The importance TE mediated regulatory rewiring

Evolutionary innovations and adaptations often require rapid and concerted changes in regulation of gene expression at many loci (Carroll 2006). It has been suggested that TEs play a key role in rewiring regulatory networks, since the dispersal of TEs that contain a regulatory element may allow for the same regulatory motif to be recruited at many genomic locations (McCLINTOCK 1951; Britten and Davidson 1969; Rebollo *et al.* 2012). A handful of recent studies have implicated TEs as drivers of key evolutionary innovations, including placentation in mammals (Lynch *et al.* 2011), or rewiring the core regulatory network of human embryonic stem cells (Kunarso *et al.* 2010). While these studies demonstrate that TEs can, in principle, contribute to the creation or rewiring of regulatory networks, they do not address the question of how often regulatory elements evolve by TE insertions versus by other mutations. That is, the importance of TEs in contributing to regulatory evolution is not known. Quantification of the role of TEs would require *a priori* knowledge of how and when regulatory networks evolve, and a detailed molecular understanding of which genes are being drawn into a regulatory network, and how. As discussed above, all these variables are understood for dosage compensation in flies.

Our previous work in *D. miranda* has shown that a helitron TE was recruited into the dosage compensation network at two independent time points. The younger 1.5 MY old neo-X chromosome of *D. miranda* is in the process of evolving dosage compensation, and dozens of new CES on this chromosome were created by insertions of the ISX element (Ellison and Bachtrog 2013). We showed that the domesticated ISX transposable element gained a novel MRE motif by a 10-bp deletion of the highly abundant ISY element in the *D. miranda* genome (Ellison and Bachtrog 2013). We also found the remnants of a related (but different) TE at CES on the older neo-X of this species (which formed roughly 13-15MY ago), but the TE was too eroded to reconstruct its evolutionary history. Here, we identified another domesticated TE that was utilized to deliver MSL binding sites to a newly formed neo-X chromosome, but no significant TE contribution was found for MSL binding site evolution in two independent neo-X chromosomes. Our data shed light on the question of when we expect TEs to be important in regulatory evolution. For TEs to contribute to regulatory rewiring, two conditions have to be met: a regulatory element (or a progenitor sequence that can easily mutate into the required binding motif) needs to be present in the TE, and that TE needs to be active in the genome (and not yet silenced by the host machinery). TEs undergo a characteristic life cycle, in where they invade a new species (or escape the genome defense by mutation) and transpose, until they are silenced by the host genome. Once a TE is robustly repressed, it no longer can serve as a vehicle to disperse regulatory elements, so the time window where a particular TE family can be domesticated is probably short, and needs to coincide with a necessity to disperse regulatory motifs. A high TE burden does increase that chance, but at a cost: maintaining active TEs in the genome allows a rapid response to evolutionary challenges, but also creates a major source of genomic mutation, illegitimate recombination, genomic rearrangements, and genome size inflation (Chuong *et al.* 2017).

Our findings support this view of a TE trade-off. The ISY element in *D. miranda* is the most highly abundant transposon in the *D. miranda* genome, and is massively contributing to the degeneration of the neo-Y in this species (Mahajan *et al.* 2018). Indeed, our genomic analysis has revealed >20,000 novel insertions of the ISY element on the neo-Y, often within genes (Mahajan *et al.* 2018). Yet, it contained a sequence that was only one mutational step away from a functional MSL binding site (i.e a single 10-bp deletion), and domestication of this element allowed for the rapid dispersal of functional binding sites for the MSL complex along the neo-X. The domestication of the TE in *D. robusta* occurred too long ago for us to reconstruct its exact evolutionary history and the potential damage its mobilization may have caused while it was active. However, consistent with a trade-off that the host genome faces, we find that *D. robusta* has a higher TE density than its sister species and also a considerably larger genome size, yet a TE contributed to wiring 100s of genes into the dosage compensation network on its neo-X.

### What makes a binding site functional?

Perhaps surprisingly, in many instances we are unable to detect specific mutations that would generate a novel MSL binding motif. Instead, we find that functional MSL-binding sites are derived from pre-sites containing the GA-rich motif that was already present in an ancestor where the neo-X is autosomal, and where these sequences do not recruit the MSL complex. The MSL binding motif is only modestly enriched on the X chromosome compared to the autosomes (only ∼2 fold), and only a small fraction of putative binding sites are actually bound by the MSL complex (Alekseyenko *et al.* 2008). The dosage compensation machinery shares this characteristic with many other sequence-specific binding factors whose predicted target motifs are often in vast excess to the sites actually utilized. It has been speculated that other genomic aspects, such as chromatin context or the 3D organization of the genome could help to distinguish between utilized and non-utilized copies of a motif. Our finding that a large number of sites can acquire the ability to recruit the MSL complex, without any associated changes at the DNA level, supports the view that epigenetic modifications or changes to the three-dimensional architecture of the genome help to ultimately determine what putative binding sites in the genome are actually utilized (Alekseyenko *et al.* 2012; Schauer *et al.* 2017). In *D. melanogaster*, the X chromosome has a unique satellite DNA composition, and it was suggested that these repeats play a primary role in determining X identity during dosage compensation (Joshi and Meller 2017). Future investigations of changes in the chromatin level, the repeat content, and the genomic architecture of these newly formed sex chromosomes will help to resolve this outstanding question.

## Methods

### Genome sequencing, assembly, and alignment

DNA was extracted from single flies using the Qiagen PureGene Kit and two paired-end Illumina sequencing libraries (male and female) were prepared for each species. The Illumina Nextera library prep kit was used for *D. melanica* and *D. robusta* (150 bp PE reads), while the Illumina TruSeq kit (100 bp PE reads) was used for the remaining species. Genome assemblies were generated for males and females separately by first error-correcting reads using *BFC* (Li 2015) and then assembling the corrected reads using *IDBA* (Peng *et al.* 2012). A whole-genome alignment was constructed using the female assemblies for the five species studied here plus *D. virilis* using *Mercator* (Dewey 2007).

### Phylogeny

To create a whole-genome phylogeny, the *D. virilis* genome was split into 250 bp windows. Each window was extracted from the *Mercator* whole-genome alignment and windows were retained if the aligned sequence from each species contained no more than 10% of positions as gaps.

Retained windows were further filtered to ensure that each window was at least 1 kb from the closest neighboring window. These windows were concatenated to produce a multiple sequence alignment containing 1.1 million positions. The RAxML rapid bootstrapping algorithm (Stamatakis 2014) was used to produce a maximum likelihood phylogeny from this alignment.

### Chromosome assignments and sex-linkage

Chromosome assignments for *D. virilis* scaffolds were obtained from (Schaeffer *et al.* 2008). The scaffolds from each species studied here were assigned to Muller elements based on their alignment to *D. virilis* scaffolds from the *Mercator* whole-genome alignment.

To determine which Muller elements are X-linked in each species, male and female Illumina reads were aligned separately to the female genome assemblies using *bowtie2* (Langmead and Salzberg 2012) and male/female coverage ratios were calculated for each female scaffold.

Y-linked scaffolds were identified from the male assemblies using *YGS* (Carvalho and Clark 2013).

### RNA-seq and roX2 identification

For each sex, heads were removed from five flies, flash frozen in liquid nitrogen and placed into Trizol for RNA extraction. The Illumina TruSeq RNA kit was used to prepare unstranded, single-end 50 bp sequencing libraries for each sex. RNA-seq data were aligned to the female reference genome assembly using *Hisat2* (Kim *et al.* 2015) and gene models were generated from the merged male+female spliced alignments, along with normalized expression values, using *StringTie* (Pertea *et al.* 2015). Male and female RNA-seq read coverage was used to identify the location of *roX2* in *D. melanica* and *D. robusta* (see **Figure 3B**), and roX2 transcripts were extracted from the genome assemblies based on the *StringTie* gene models. The *D. nigromelanica* roX2 transcript was identified based on homology to the *D. melanica* transcript using *Exonerate* (Slater and Birney 2005).

### Species pairs and neo-X/Y chromosome divergence

For each species, *D. melanogaster* peptides were searched against the set of neo-X and Y-linked scaffolds using a translated BLAST search (Camacho *et al.* 2009). The resulting neo-X and Y-linked gene models were further refined using *Exonerate* and their coding sequence was aligned using the codon model in *PRANK* (Löytynoja 2014). Ks values were calculated for each neo-X/Y pair using *KaKs_Calculator* (Zhang *et al.* 2006). For species divergences, orthologous genes were identified using the *D. robusta* gene models from *StringTie* and the *Mercator* whole genome alignment. Refinement of gene models, alignment, and Ks values were obtained as described above.

### Divergence time estimates

We used a neutral mutation rate estimate of 3.46 x 10^-9^ per base per generation, which was experimentally determined from *D. melanogaster* (Keightley *et al.* 2009). The species studied here have a generation time that is roughly twice as long as *D. melanogaster* and we therefore used the lower bound of the estimate of the number of generations per year for Drosophilids (5 generations) (Cutter 2008) to convert the mutation rate to time-based units (1.73 x 10^-8^ mutations per base per year).

### Chromatin isolation by RNA purification (ChIRP)

ChIRP sequencing libraries were prepared according to the published protocol (Chu *et al.* 2012), using the Drosophila-specific modifications described in (Quinn *et al.* 2016). For each species, ChIRP libraries were prepared from 2 different pools of 6 probes (**Table S3**), which were tiled across the roX2 transcript. Input control libraries were also prepared for each species by extracting DNA from an aliquot of the cell lysate immediately prior to probe hybridization. Wandering third instar larvae were used for *D. melanica* and *D. robusta*. Due to difficulty in collecting sufficient larvae for *D. nigromelanica*, adult males were used instead. 100 bp paired-end Illumina reads were generated for each pool, for each species, and aligned to the female reference genome assembly using *bowtie2*. Peaks of roX2 binding were identified by running *MACS* (Zhang *et al.* 2008) on each pool separately, along with the control library, and a final set of peaks were generated by retaining only the subset of peaks that were identified in both pools.

### Identification of Chromatin Entry Sites (CES)

The ChIRP libraries varied in overall signal versus background, likely due to differences in the hybridization efficiency of the different probesets. For each species, we calculated the average fold-enrichment (treatment versus control) across all peaks as a measure of overall ChIRP signal. The *D. robusta* libraries showed the highest signal and we used the same enrichment threshold (20) that was previously used to identify CES from roX2 peaks (Quinn *et al.* 2016). For the remaining species, we identified CES by scaling the enrichment threshold in proportion to our measure of ChIRP signal.

### Motif Identification and Evolution

We defined the location of chromatin entry sites as a 500 bp region centered on the summit of the roX2-bound peak. We extracted the sequence from these regions, for the ancestral and neo-X chromosomes separately, and used the ZOOPS model in *MEME* (Bailey *et al.* 2009) to identify enriched sequence motifs. We used *FIMO* (Bailey *et al.* 2009) to determine the location of MRE and pion-X motifs within each CES and assigned each CES as containing either a MRE motif or pion-X motif (whichever match had a higher score), or no motif (if there was no match to either motif). We used the *Mercator* whole-genome alignment to assess orthology of CES as well as individual motifs. For each neo-X CES, we manually viewed the alignment of its sequence motif with the orthologous sequences from the other five species to determine the mutational mechanism that gave rise to the motif.

### Transposable element (TE) identification

De novo TE identification was performed for each species using *RepeatModeler* (https://github.com/rmhubley/RepeatModeler). To identify the genomic locations of TE families, *RepeatMasker* (https://github.com/rmhubley/RepeatMasker) was used with the *RepeatModeler* consensus sequences as the repeat library. *D. robusta* TEs that overlapped chromatin entry sites were further classified using *CENSOR* (Kohany *et al.* 2006) and *RepBase* (Bao *et al.* 2015).

### Repeat Density Estimates

For each species, we used *RepeatMasker* to separately identify simple repeats and transposable elements. Because the percentage of the genome assembly that falls into these two categories will be affected by differences in total assembly size between species, we used an alternative approach for determining the density of these repeat classes. For each species, we permuted the location of 1000 1Kb windows for 1000 permutations. For each iteration we determined the number of windows that overlapped a simple repeat and the number of windows that overlapped a transposable element, which we termed the “repeat density metric”.

## Acknowledgements

Funded by NIH grants (R01GM076007, GM101255 and R01GM093182) to DB. We thank J. Quinn for advise on ChIRPs.

## Author contributions

CE generated all the genomic data, RNA-seq and ChIRP data, and performed the bioinformatics analysis. D.B. oversaw the project and wrote the manuscript with input from all authors.

## Supplementary data

**Figure S1.**
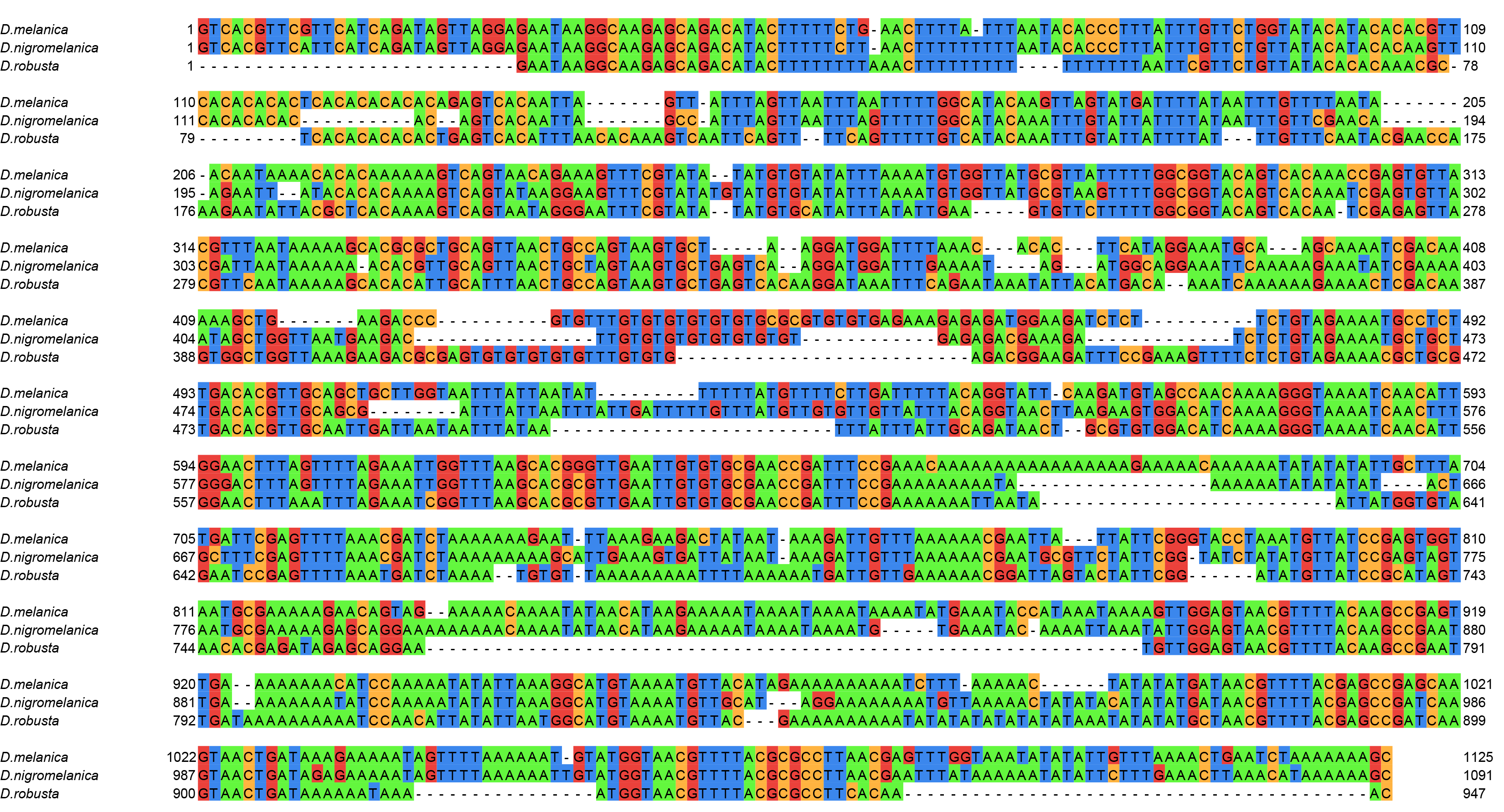
We identified the roX2 transcript in *D. melanica* and *D. robusta* by using gene synteny and RNA-seq data. The *D. nigromelanica* roX2 transcript was identified based on sequence homology to the transcripts from *D. melanica* and *robusta*. A multiple sequence alignment of the three transcripts is shown here.

**Figure S2.**
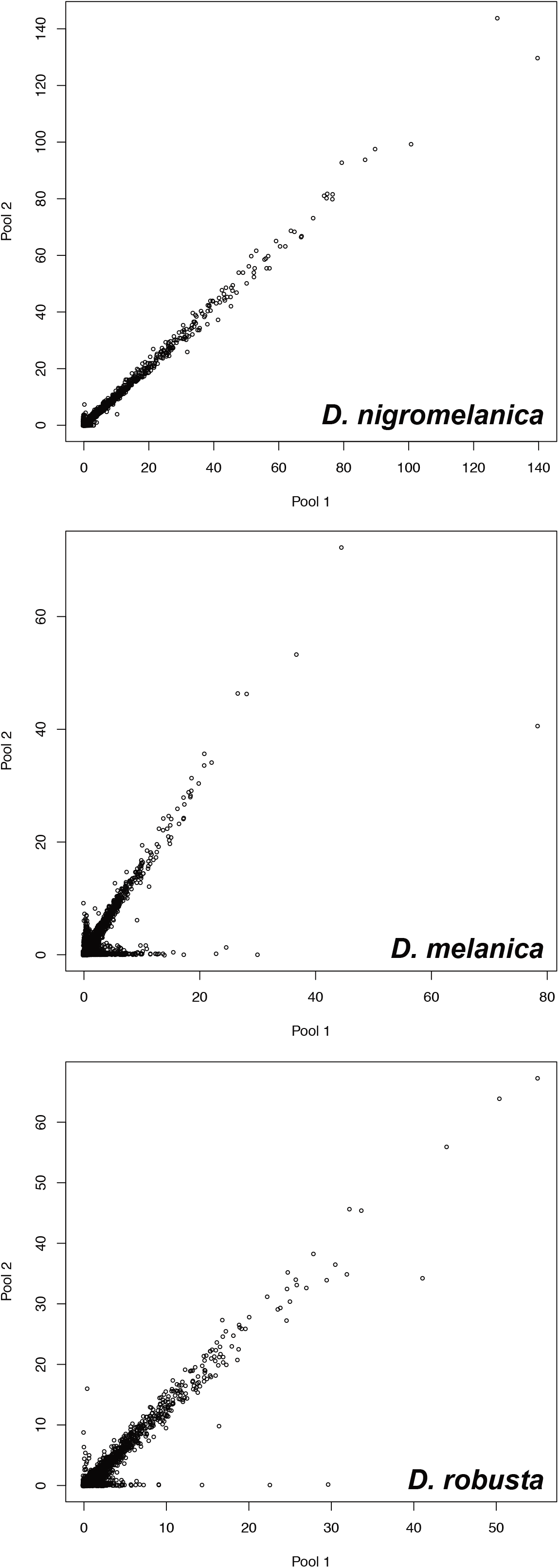
For each species, two ChIRP-seq libraries were prepared using two pools of non-overlapping probes tiled across the roX2 transcript. Each dot in the scatterplot represents a roX2-bound peak identified in one or both pools, and its location in the plot reflects the fold-enrichment (ChIRP/input control) of that region in each pool.

**Figure S3.**
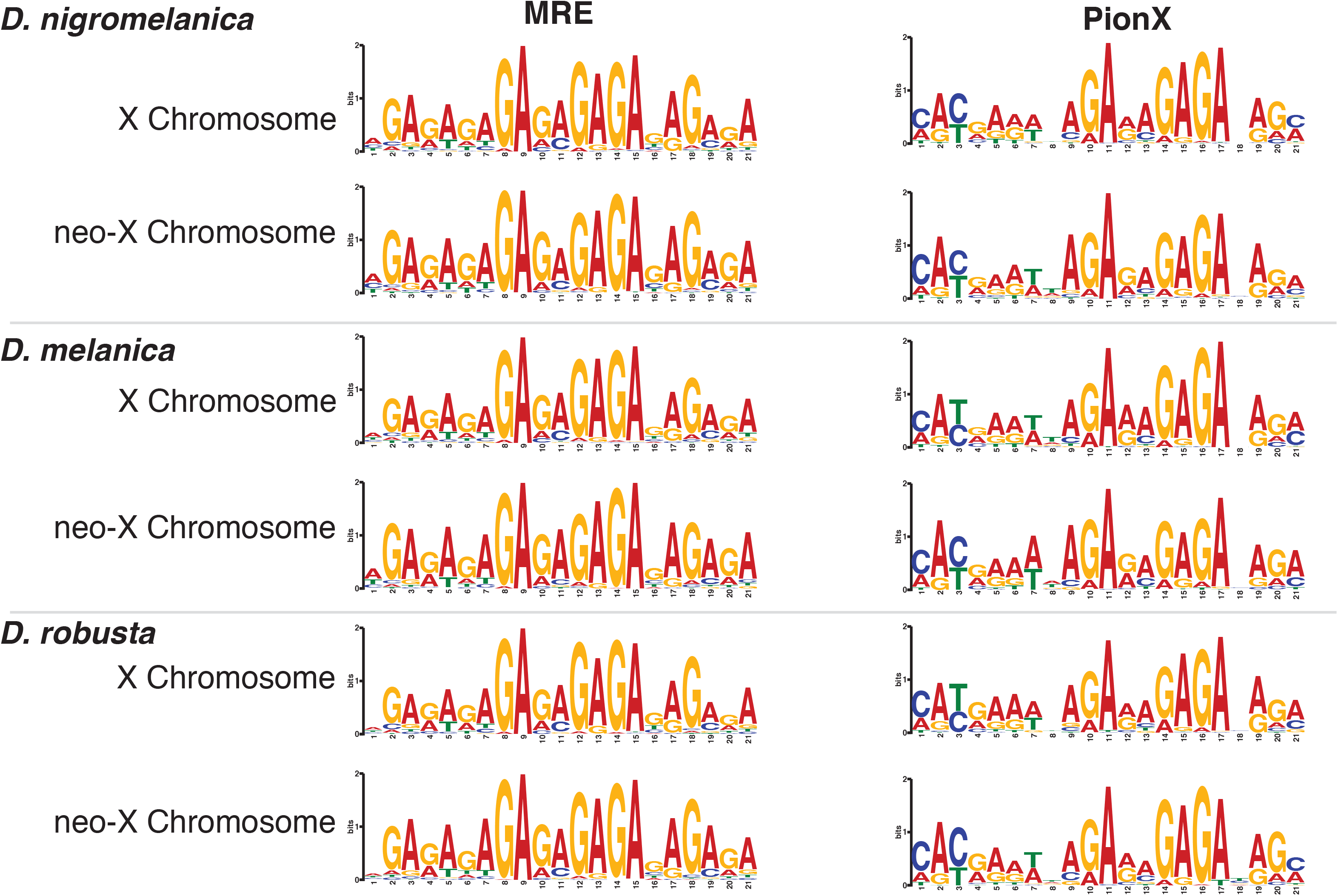
We assigned each CES as containing either an MRE motif, a pion-X motif, or no motif, based on searching the consensus sequences of these motifs against the CES sequences and identifying the best-scoring match. To validate these matches, we extracted the MRE or pion-X element from each assigned CES sequence and created a consensus logo for each X chromosome of each species. The consensus motifs are very similar to the MRE and pion-X consensus motifs discovered in *D. melanogaster*.

**Figure S4.**
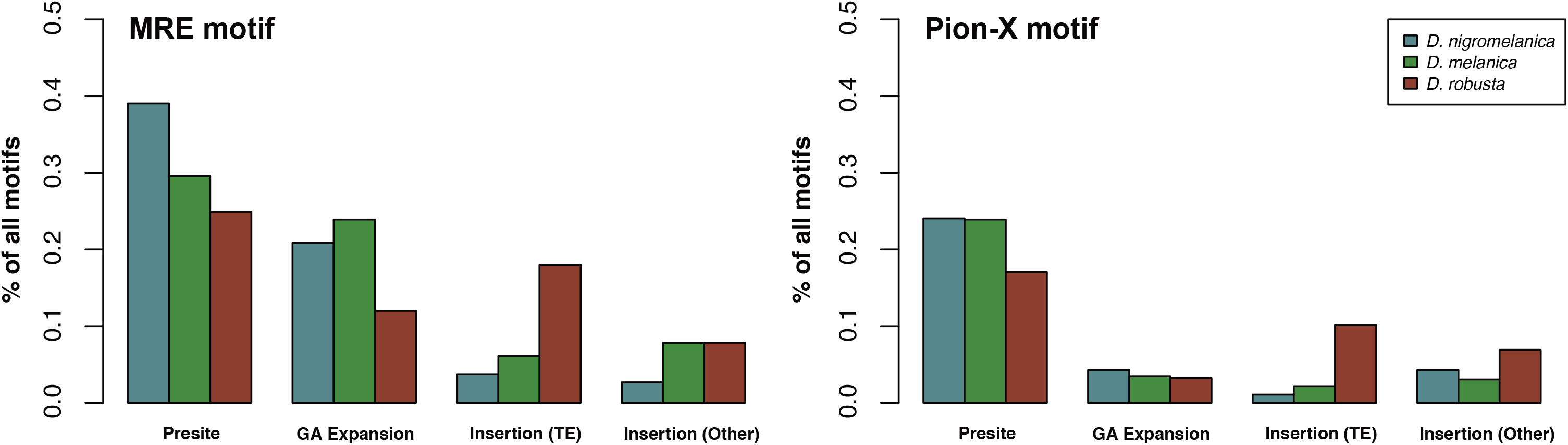
Evolutionary mechanism that gave rise to the MRE or pion-X motifs found within each CES on the neo-X chromosome of each species.

**Figure S5.**
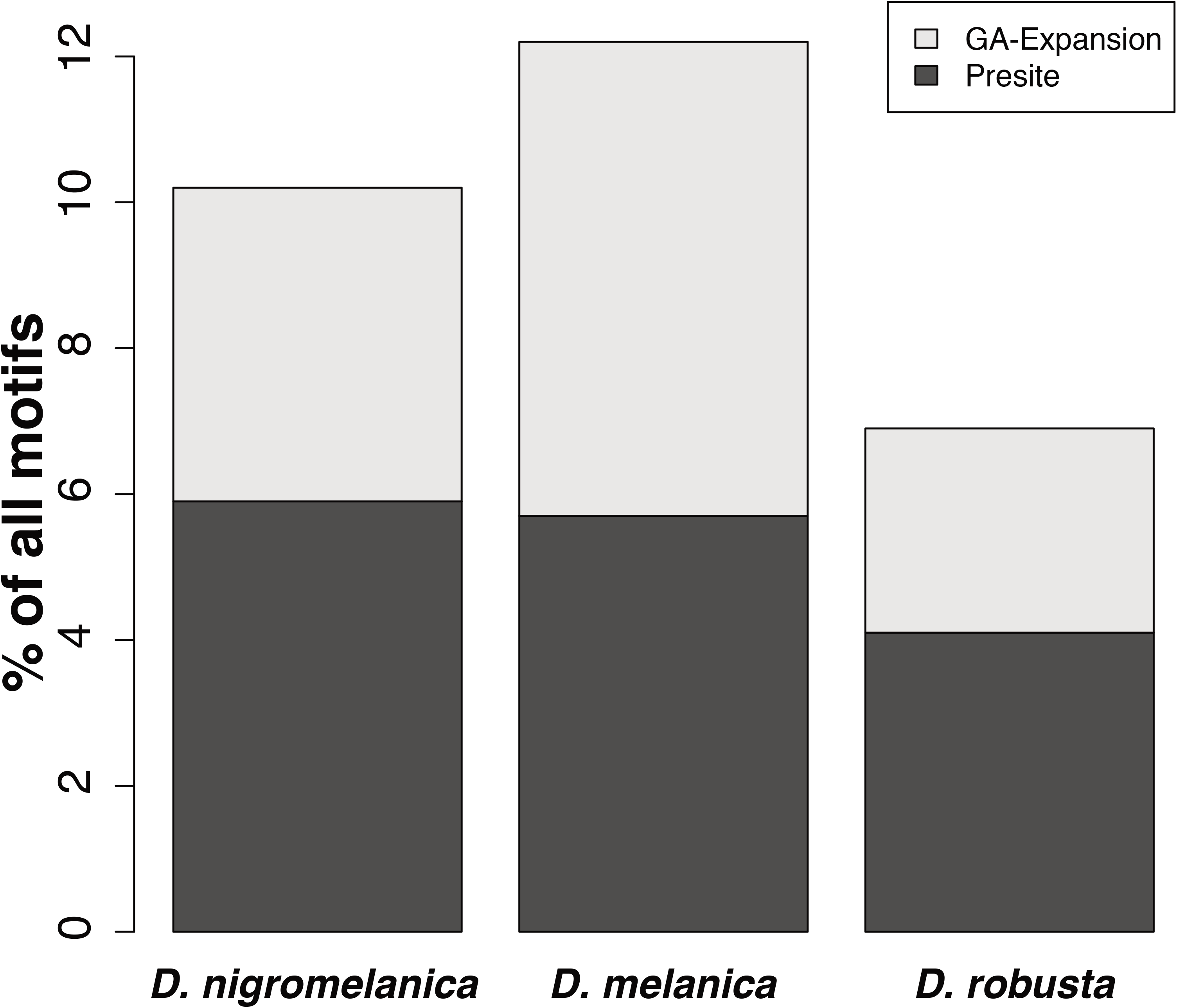
For each species, we identified both MRE and pion-X motifs in the 3’ 100 bp of introns, where the GA-rich polypyrimidine tracts involved in RNA splicing are located. Shown here are combined counts for MRE and pion-X motifs. Less than 15% of all motifs were derived from these tracts in any species. For those that did arise from polypyrimidine tracts, about half served as a ready-made motif (i.e. presite) whereas in the other half, additional GA-expansion mutations occurred at the tract location.

**Table S1.** Genomic data generated.

| Stock Center | Stock Number | Genus | Species | Assembly | Karyotype |
| --- | --- | --- | --- | --- | --- |
| UCSD | 15020-1111.10 | *Drosophila* | *robusta* | 184 Mb | A + D |
| UCSD | 15030-1151.01 | *Drosophila* | *micromelanica* | 146 Mb | A |
| UCSD | 15030-1141.03 | *Drosophila* | *melanica* | 150 Mb | A + D |
| EHIME | E-22901 | *Drosophila* | *nigromelanica* | 164 Mb | A + C |
| EHIME | E-14007 | *Drosophila* | *lacertosa* | 155 Mb | A + D |

**Table S2.** Inferred protein divergence of homologous gene pairs identified on the neo-sex chromosomes of *D. nigromelanica*, *D. melanica, D. lacertosa* and *D. robusta*.

|  | No. of genes | Ka | Ks |
| --- | --- | --- | --- |
| *D. nigromelanica* | 118 | 0.0579 | 0.1591 |
| *D. melanica* | 20 | 0.0430 | 0.2590 |
| *D. robusta* | 16 | 0.0661 | 0.3948 |
| *D. lacertosa* | 11 | 0.0715 | 0.5223 |

**Table S3.**
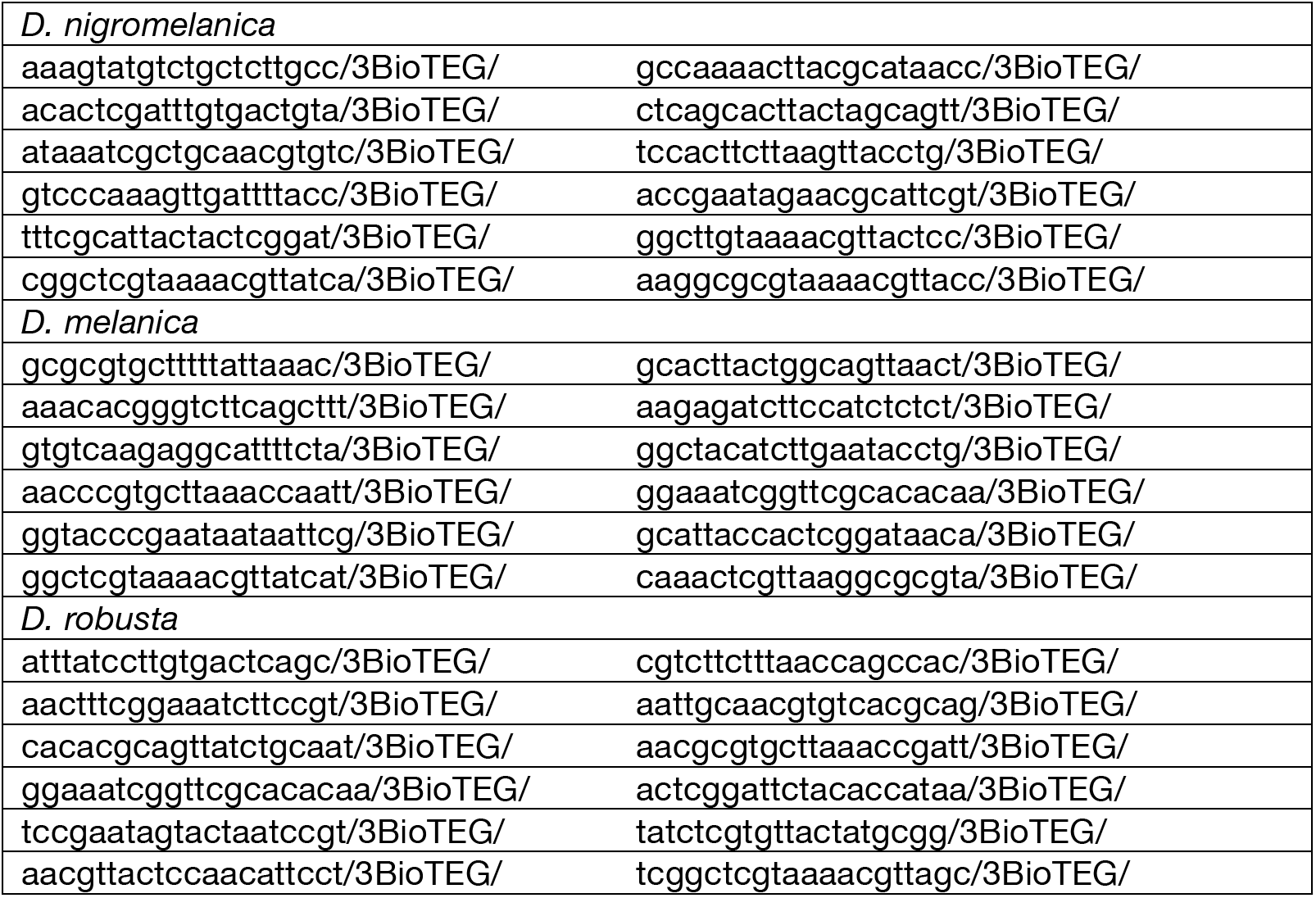
Oligos used for *roX2* ChIRP in *D. nigromelanica*, *D. melanica* and *D. robusta*.

**Table S4.**
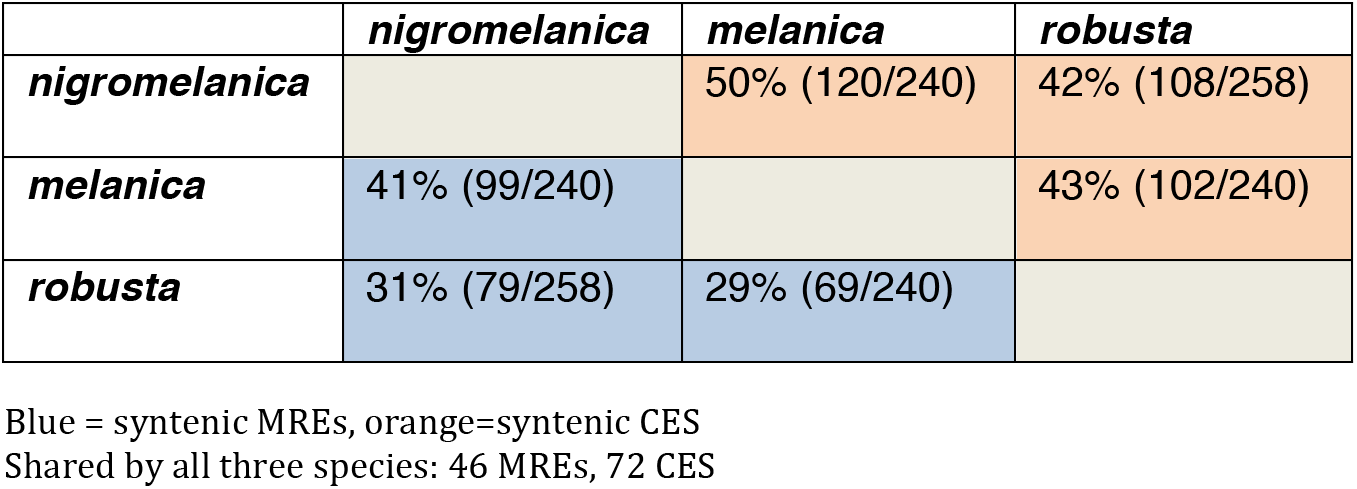
Synteny of CES on the ancestral X chromosome across species.

**Table S5.** Transposable elements contributing to CES evolution in *D. robusta*.

| RepeatModeler ID | RepBase Match | MRE | PION-X | TOTAL |
| --- | --- | --- | --- | --- |
| rnd-5_family-1861* | Galileo_DM | 17 | 7 | 24 |
| rnd-5_family-495 | GA-rich_Satellite/Low-complexity | 8 | 0 | 8 |
| rnd-5_family-25933,<br>rnd-5_family-1154* | Galileo_DM | 7 | 1 | 8 |
| rnd-5_family-3008 | Unknown | 0 | 3 | 3 |
| rnd-5_family-6205 | Unknown | 0 | 2 | 2 |
| rnd-2_family-56 | Helitron-2N1_DVir | 2 | 0 | 2 |
| rnd-5_family-9219 | Unknown | 1 | 0 | 1 |
| rnd-5_family-6299 | Helitron-2_DVir | 1 | 1 | 2 |
| rnd-5_family-5467 | MarN-2_DEu | 0 | 1 | 1 |
| rnd-5_family-5050 | Unknown | 0 | 1 | 1 |
| rnd-5_family-1890 | Satellite/Low-complexity | 1 | 0 | 1 |
| rnd-5_family-142 | Loa-2_Dper | 1 | 0 | 1 |
| rnd-5_family-1163 | Gypsy-4_DGri-I | 1 | 0 | 1 |
| rnd-4_family-95 | Unknown | 0 | 1 | 1 |
| rnd-3_family-58<br>rnd-4_family-310<br>rnd-5_family-1710 | PERI_helitron_Dbuz | 0 | 3 | 3 |
| rnd-4_family-94 | Helitron1_Dmoj | 0 | 1 | 1 |
| rnd-4_family-1707 | Helitron-1_DVir | 0 | 1 | 1 |

\*These elements show only short homology to different regions of Galileo\_DM and are not homologous to each other, nor are they subparts of a single larger element

